# An evolutionary epigenetic clock in plants

**DOI:** 10.1101/2023.03.15.532766

**Authors:** N Yao, Z Zhang, L Yu, R Hazarika, C Yu, H Jang, LM Smith, J Ton, L Liu, J Stachowicz, TBH Reusch, RJ Schmitz, F Johannes

## Abstract

Molecular clocks are the basis for dating the divergence between lineages over macro-evolutionary timescales (~10^5^-10^8^ years). However, classical DNA-based clocks tick too slowly to inform us about the recent past. Here, we demonstrate that stochastic DNA methylation changes at a subset of cytosines in plant genomes possess a clock-like behavior. This ‘epimutation-clock’ is orders of magnitude faster than DNA-based clocks and enables phylogenetic explorations on a scale of years to centuries. We show experimentally that epimutation-clocks recapitulate known topologies and branching times of intra-species phylogenetic trees in the selfing plant *A. thaliana* and the clonal seagrass *Z. marina*, which represent two major modes of plant reproduction. This discovery will open new possibilities for high-resolution temporal studies of plant biodiversity.

---

Reconstructing the tree-of-life is a central goal in evolutionary biology. A key challenge is to infer the approximate date when two lineages diverged from each other in the past (*1*, *2*). In addition to fossil and archaeological evidence, molecular clocks have emerged as an important tool to perform such dating (*2*, *3*). Constant-rate clock calibrations such as those originally introduced by Zuckerkandl and Pauling (*4*) are based on the premise that neutral mutations in DNA (or proteins) accumulate at a fixed rate, so that nucleotide differences increase with time. If the mutation rate is known, it becomes possible to deduce when two lineages shared their most recent common ancestor (MRCA). Although the modern use of molecular clocks relies on a number of strong modeling assumptions (*5*), in practice, they are often the only means to obtain temporal information for parts of a phylogeny where fossil or archaeological records are lacking (*3*).

The low mutation rate found in most species limits the use of DNA-based clocks. With a rate of ~10^-9^-10^-8^ (per site per year), they may offer sufficient temporal resolution over larger timescales (~10^4^ to 10^8^ years) but are less accurate in recent time (< 10^3^ years from the present), as too few mutations accumulate to permit reliable tree inference and dating (*6*). However, it may be of interest to infer shallow divergence times of just a few decades to hundreds of years, for example, when assessing associations with species range shifts, colonization events, or environmental changes (*7*). In self-fertilizing or clonal species with short life cycles, new lineages can diverge rapidly due to extensive genetic drift, restricted gene flow, and divergent natural selection (*8*). In the recent past, many such events have co-occurred with the emergence of modern civilizations and may in part even be driven by human activities (e.g., migration or trade). To improve the resolution in studying shallow divergences and their timing, a new class of molecular clocks is needed, whose tick-rate is orders of magnitude faster than that of DNA. We recently proposed that DNA cytosine methylation could provide a biomolecular basis for such a clock in plants (*9*), but this possibility has not been explored rigorously.

DNA cytosine methylation is a conserved base modification in eukaryotes (*10*). Stochastic enzymatic failure or off-target DNA methyltransferase activity at CG dinucleotides leads to lasting methylation losses or gains (i.e., epimutations) in daughter cells and their decedent cell lineages (*11*). Such CG methylation changes have been observed within the lifetime of mammals and have been extensively exploited as a DNA methylation clock of aging (*12*). However, unlike in mammals, many somatically-acquired CG epimutations are stably inherited across clonal and sexual generations in plants (*13*–*15*), and thus hold high-resolution information about the evolutionary histories of cell lines or clonal and sibling lineages (*11*, *16*). Estimates in several plant species indicate that CG epimutations are effectively neutral at the genome-wide scale and occur at a rate that is ~10,000 - 100,000 times higher than the genetic mutation rate per unit time (*13*, *14*, *17*–*20*). Here we show that the rapid accumulation of CG epimutations in plant genomes defines a fast-ticking evolutionary clock (henceforth ‘epimutation-clock’), which can be used for the reconstruction and dating of phylogenies.

We set out to construct a robust epimutation-clock by first searching for genomic regions whose CG epimutation rates are invariant to genetic and environmental perturbations. To do this, we used the selfing plant *Arabidopsis thaliana* as a model to generate mutation accumulation lines (MA-lines) from seven diverse natural accessions (i.e., genotypes) as founders (**Fig. 1A, Fig. S1A, Materials and Methods**). These MA pedigrees had a depth of 17 generations with nine whole genome bisulfite sequencing (WGBS) measurements per pedigree, on average (see **Fig. S1A** for details). To evaluate the impact of environmental factors, we also generated *A. thaliana* MA lines grown under biotic stress (repeated exposure to *Pseudomonas syringae* and salicylic acid) and combined these data with published MA-lines grown under abiotic stress (exposure to high salinity and drought conditions, (*21*, *22*), **Fig. 1A, Fig. S1B-D, Materials and Methods**). The depth of these latter MA pedigrees varied from 7-13 generations and had ~9 WGBS measurements per pedigree.

**Fig. 1.**
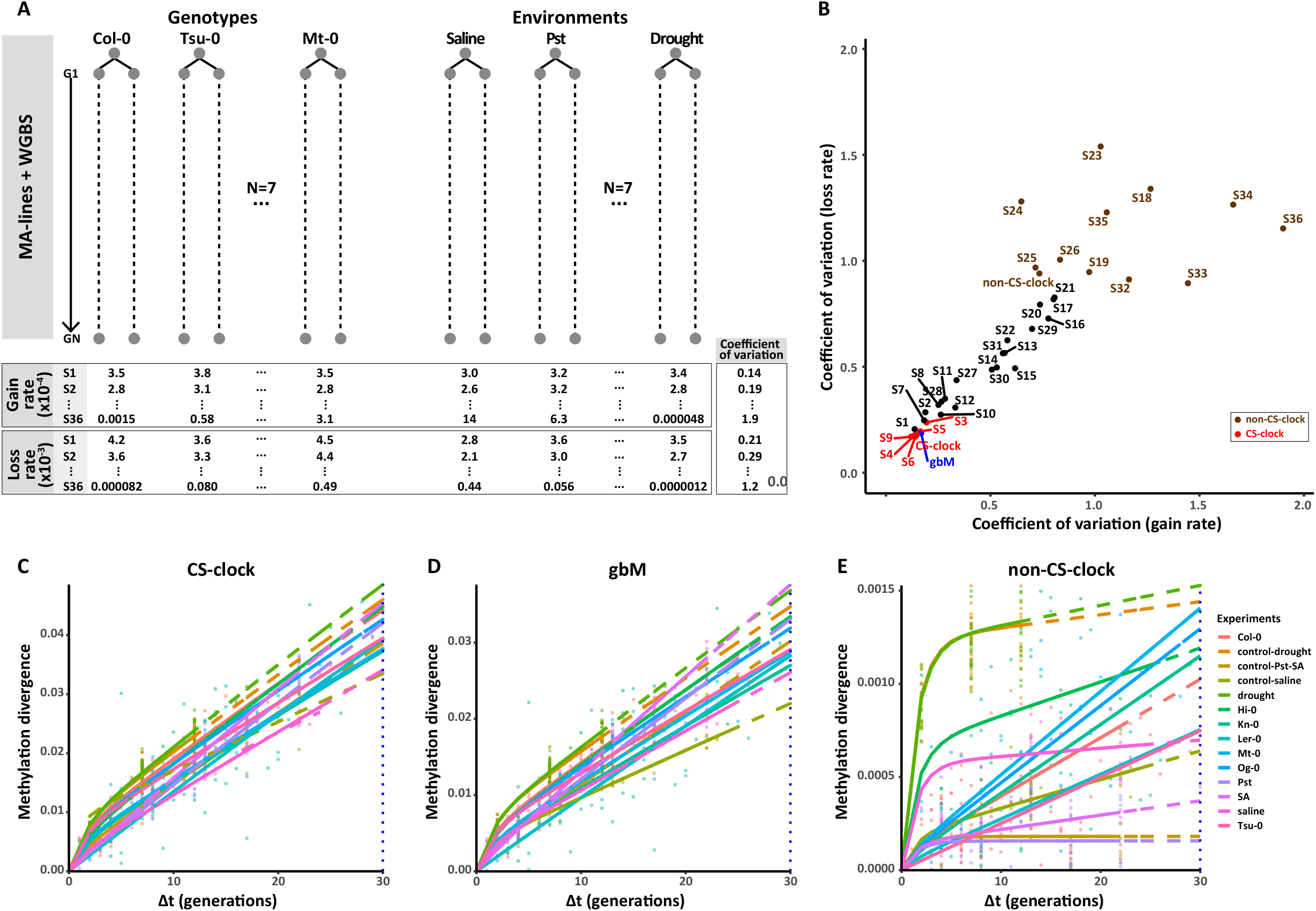
Discovery of epigenetic clock regions. **(A)** We sought to identify genomic regions whose CG epimutation rates are invariable across genetic and environmental perturbations. To that end, we created seven MA pedigrees in various genetic backgrounds by using different natural accessions as founders (Col-0, Hi-0, Kn-0, Ler-0, Mt-0, Og-0, and Tsu-0). In addition, we generated MA lines grown under biotic stress (repeated exposure to *Pseudomonas syringae* and salicylic acid) and combined these data with published MA-lines grown under abiotic stress (exposure to high salinity and drought conditions). Using multigenerational WGBS data from these MA lines, we estimated CG epimutation rates in each of 36 annotated Arabidopsis chromatin states (CS). Detailed information about pedigree topologies and WGBS sampling can be found in Fig. S1. For each CS, the coefficient of variation (CV) in the gain and loss rates were calculated across pedigrees. **(B)** Low CV for the gain and the loss rates were identified in a cluster of CS including S3, S4, S5, S6, and S9. Genomic regions indexed by these CS were defined as epigenetic clock regions (CS-clock). For comparison, a cluster of CS including S18, S19, S23, S24, S25, S26, S32, S33, S34, S35, and S36 displayed high CVs, which were defined as non-clock regions (non-CS-clock). **(C-E)** CG epimutation accumulation is plotted as a function of divergence time (Δt) for each MA pedigrees. For CS-clock regions (D), epimutation accumulation patterns were much more invariable across pedigrees than for non-CS-clock regions (E). Epimutation accumulation in gbM genes (D) closely resembled CS-clock regions, indicating that gbM genes can serve as a proxy for an epigenetic clock.

Epimutation analysis of these different MA-lines revealed specific genomic regions, whose CG epimutation rates were largely invariable across the 14 different genetic or environmental perturbation experiments (**Fig. 1B-E, Fig. S2, Table S1, Materials and Methods**). These clock-like regions were in stark contrast with other regions where rates were 5-fold more variable on average (**Fig. 1B, Table S1**). We found that clock-like regions displayed specific epigenomic features and comprised about 16.1% of all CGs in the genome (~896,323 of CG total sites, **Fig. 1B**). The majority of clock-like regions (~60%) were located within gene body methylated (gbM) genes. In *A. thaliana*, gbM genes comprise a subclass of ~5,000 genes that display elevated CG methylation (*23*). Although the precise function of gbM is unclear, the nucleotide sequences and steady-state methylation levels of these genes are generally conserved across diverse plant species (*24*, *25*). Interestingly, we recently identified these same regions as epimutation hotspots in *A. thaliana* (*26*). Their average epimutation rates exceed the genome-wide average by one order of magnitude (~10^-3^ versus 10^-4^ per CG site, per haploid genome, per generation). Hence, the biomolecular properties of these regions form the basis for a robust epimutation-clock, whose fast substitution rate can facilitate high-resolution inference about divergence events in the recent past. We here use the term “mCG substitution rate” to refer to the number of fixed CG methylation changes that occur over a unit period of time.

To confirm the existence of such clock-like regions, we analyzed the largest *A. thaliana* MA pedigree available (here named MA1_1 and MA1_2, see **Fig. 2A**) (*13*, *14*). This MA pedigree features 15 independent lineages with a maximum depth of 32 generations. WGBS samples were available from generations 3, 31, and 32. In total, we detected 46,597 segregating CG epimutations within the clock-like regions after ~31 generations. By contrast, only 99 segregating SNPs were detected genome-wide over this timescale (*27*). This shows that epimutations in the relatively small clock-like regions are overall much more abundant than genome-wide SNPs. Using pairwise distances based on the GTR2 substitution model (**Materials and Methods**), we performed neighbor-joining clustering of the samples on the basis of their mCG status within the clock-like regions and were able to recapitulate the known topology (**Fig. 2B**). Hence, the rapid accumulation of epimutations is highly informative about divergence events as recent as a few generations.

**Fig. 2.**
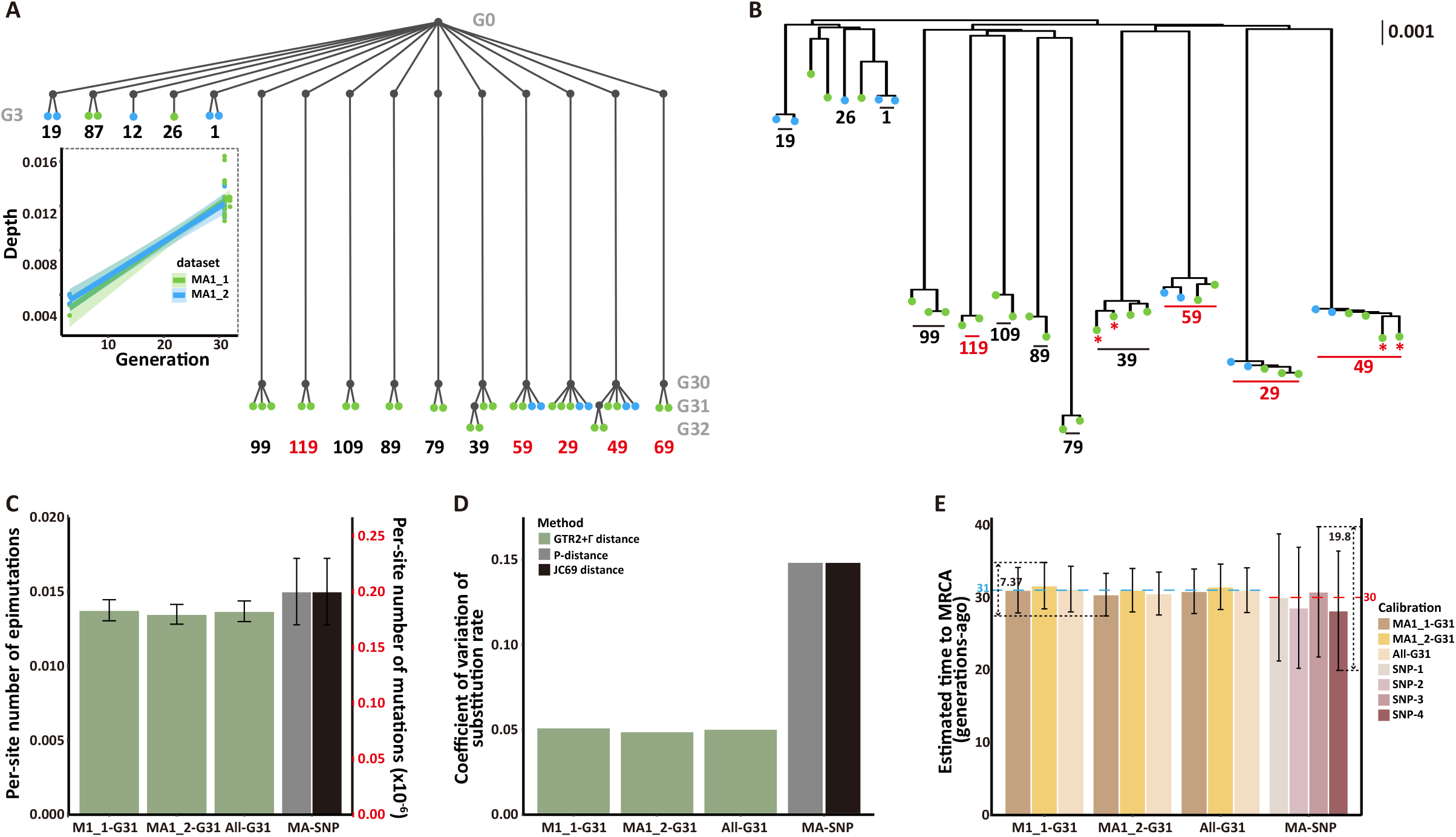
Evolutionary histories of *A. thaliana* MA lines. **(A)** The MA lines shared a common ancestor and were maintained by single-seed descent for 3, 31, and 32 generations (G3, G31, and G32). Their DNA methylomes were sequenced by Becker et al. (MA1_1, green dots) and Schmitz et al. (MA1_2, blue dots) separately. In addition, Ossowski et al. sequenced the genomes of five G30 individuals from 29, 49, 59, 69, and 119 (orange, 27). **(B)** We inferred the phylogeny of these MA lines using our epimutation clock. The inferred topology recapitulates the known evolutionary relationships among the sequenced samples. The G32 individuals are marked with a red *. **(C)** We estimated the number of accumulated substitutions on each lineage from the phylogeny (i.e., depth of tip node, Materials and Methods). The error bars represent the standard error of accumulated substitutions per lineage. The average depth of G31 individuals is consistent with the clock-like accumulation of epimutations in the different MA lineages. For comparison, the number of SNPs per G30 lineage from the previous study (27) is also shown. **(D)** The coefficient of variation (CV) of the estimated substitution rate. The CV of the substitution rate from segregating SNPs is significantly higher than the CV of the substitution rate from the epimutation clock. **(E)** With the estimated substitution rates, we inferred the time to the most recent common ancestor (MRCA). The error bars show 95% confidence intervals. The blue line indicates 31 generations. The red dash line indicates 30 generations. The epimutation clock not only estimates the correct time to the MRCA but also shows higher consistency than the times estimated with segregating SNPs. The SNP-1 is the mean of SNP substitution rates that was calibrated from the SNP phylogeny of the MA lines. SNP-2, SNP-3, and SNP-4 are four published estimates of SNP substitution rates (27, 31)

Estimates of the mCG substitution rates were highly consistent across lineages (**Fig. 2C-D**), yielding rates of 4.43 ± 0.229 x 10^-4^ and 4.34 ± 0.214 x 10^-4^ (± SE, **Supplementary text**, **Table S3**) for MA1_1-G31 and MA1_2-G31, respectively. We then applied the mean mCG substitution rate of MA1_1-G31 to the methylation data of MA1_2-G31 to infer the time until its MRCA. This estimate indicated that the MRCA lived approximately 30.4 ± 2.94 generations ago (95% CI, **Fig 2E, Table S4**), which is remarkably close to the actual depth of the pedigree (31 generations). By contrast, attempts to date the MRCA using available SNP data from the MA lines were biased and more variable, yielding an estimate of 28.2 ± 8.22 generations (95% CI), an uncertainty of nearly ~29.19% of the total age of the phylogeny (**Fig. 2E, Table S4, Materials and Methods**). Together these results indicate that CG epimutations are much more robust and informative over these short timescales than DNA mutations, a conclusion that is strongly supported by theoretical arguments and extensive simulation studies ((*6*, *28*), **Table S5**).

We sought to extrapolate these insights to natural settings. Hagmann et al. (2015) sequenced the genomes and DNA methylomes of 13 *A. thaliana* accessions collected around the Great Lakes and the East Coast of the United States (**Fig. 3A**), which all belong to a large haplogroup HPG1 (*29*, *30*). As *A. thaliana* is not native to North America, it has been hypothesized that these lineages were introduced with the arrival of the early European settlers (*30*, *31*). The average genetic distance between the 13 lineages and the HPG1 pseudo reference genome is about 245 nucleotide positions, providing evidence for their close kinship. In line with previous work (*30*, *32*), clustering accessions based on CG methylation produced nearly identical phylogenetic trees compared to SNP-based clustering, which indicates that they can capture the same evolutionary relationships between lineages (**Fig. 3B-C**). Attempts to date the MRCA for 12 non-recombinant lineages (*30*, *31*) based on the available SNP data and published SNP substitution rates from mutation accumulation experiments (*27*, *31*) yielded estimates ranging from the year 1792 ± 59.2 years (95% CI) to 1766 ± 66.2 years (95% CI) (**Fig 3F**, **Table S7**). As an alternative method, we also combined SNP data from these 12 taxa (ingroups) with that of different herbarium samples (outgroups) as input for Baysian tip calibration with BEAST (*33*). The mean values of the estimated divergence time of the 12 taxa ranged from the year ~1949 to the year ~1792, depending on the set of herbarium outgroups used (**Fig. 3G, Table S8, Supplementary text**). This extensive uncertainty is partly corroborated by our simulation studies, which indicate that a DNA mutation rate of 10^-9^ would yield large estimation errors for founding events occurring as recently as a few centuries ago (**Table S5**, **Supplementary text**).

**Fig.3.**
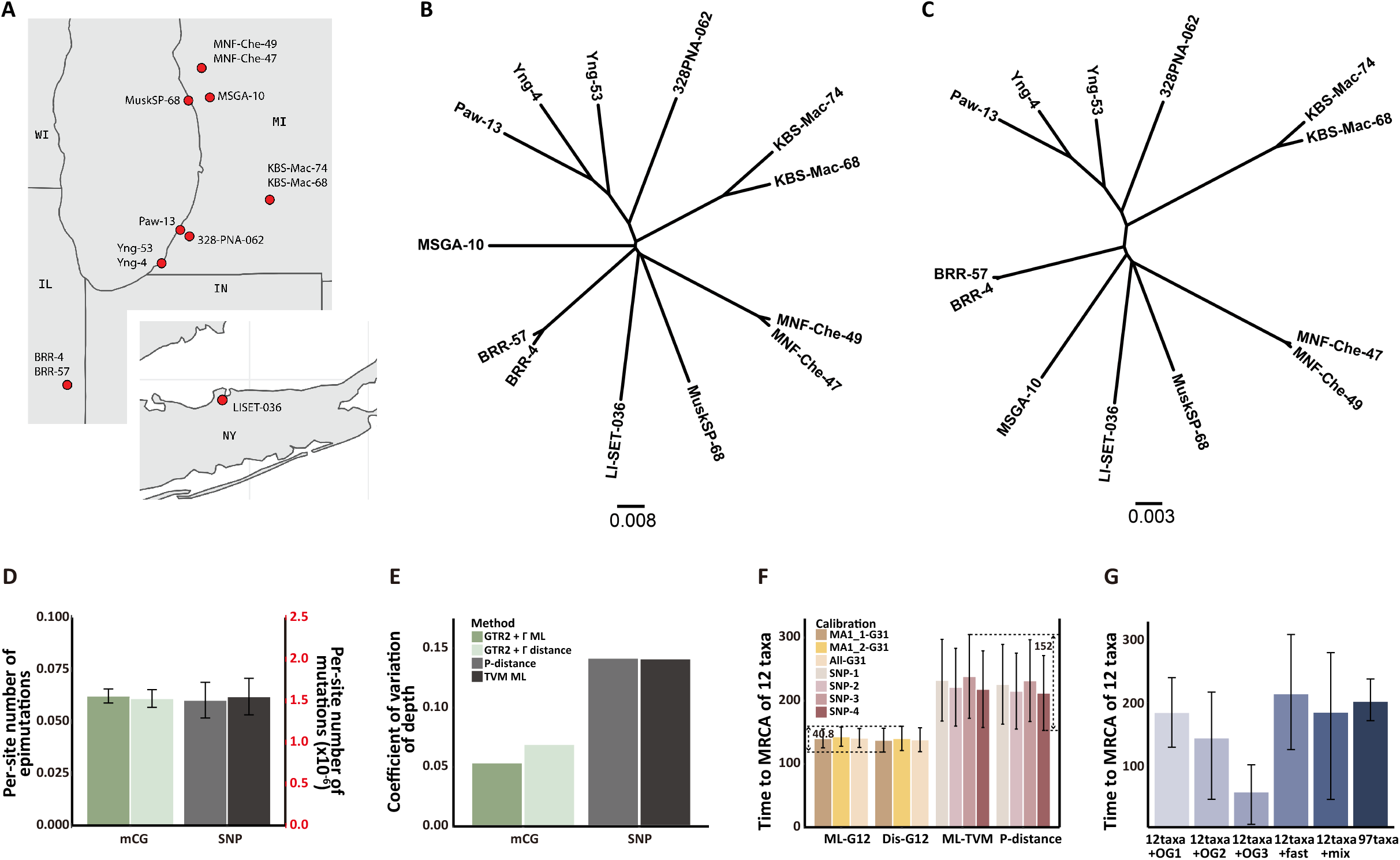
Evolutionary histories of *A. thaliana* North American accessions. **(A)** We re-analyzed the DNA methylomes of 13 samples from (30) and used them for phylogenetic inference. The selected samples all belong to a single haplogroup (HPG1, 29, 30). **(B)** The phylogeny was inferred with the same epimutation clock regions that were used in MA1_1 and MA1_2. **(C)** We reanalyzed the published set of segregating SNPs to generate the phylogeny of North American accessions (30). **(D)** The number of substitutions per lineage (± standard error). We removed Yng-53, a hybrid taxon (30, 31), and estimated the number of substitutions on the rest of the 12 lineages. **(E)** The coefficient of variation of depth suggests the SNP-based estimation has higher uncertainty than the estimation based on epimutation clock regions. **(F)** The estimated times between a modern taxon and the most recent common ancestor (MRCA) of 12 taxa. The error bars indicate 95% confidence intervals. All substitution rates of the epigenetic clock, which were calibrated from different data sources, indicate that the MRCA lived in 1860 ± 20 years. However, using four different published substitution rates of SNPs from (30) shows the MRCA lived between 1700 and 1840 (Materials and Methods). **(G)** Dating divergence event of 12 taxa with Bayesian tip calibration. We re-analyzed segregating SNPs from a total of 70 modern and 27 herbarium specimen samples from a previous study (31). In the phylogeny with all 97 samples, the MRCA of 12 taxa lived between 1769 and 1835 (95% high posterior density interval (HPD), mean=202 years ago). However, the phylogenies with only the 12 modern taxa and a part of herbarium specimen samples produced larger HPD intervals. The mean values of tMRCA varied with the sample collection dates and substitution rates on the lineages. Outgroups “OG1”, “OG2”, and “OG3” are specimens that were collected in 19 century, 1900-1950, and 1950-1985. The outgroup “fast” includes 6 fast-evolving taxa, and the outgroup “mix” has extra 3 slow-evolving taxa.

To help resolve the temporal questions about the origin of these lineages, we reanalyzed the WGBS data and applied the experimentally calibrated epimutation-clock. Our analysis placed the MRCA of these 12 lineages in the year 1863 ± 19 years (95% CI, **Fig. 3D-F, Table S7**). This estimate provides solid evidence for a very recent MRCA consistent with a previous study using an independent method (*30*). As North American herbarium samples of *A. thaliana* can be found that are older than this date (*31*), our results support the notion that these lineages belong to a clade of a larger phylogeny whose breadth has not been sufficiently sampled and/or are the result of serial founding events on this continent. Broader sampling of North American accessions would be necessary to further study the introducing time of *A. thaliana*.

The rapid formation and radiation of new plant lineages are particularly prevalent in species with facultative clonal reproduction through the production of runners, stolons or tillers. These comprise an estimated 40% of plant species on earth (*34*), many of which are of significant ecological relevance. The marine flowering plant *Zostera marina* is an important example of these. As most seagrasses, *Z. marina* is the foundation of entire ecosystems (*35*). It has important ecosystem functions in carbon storage, biodiversity enhancement, coastal protection, and is currently being developed as a seagrass genomic model (*36*). Along with sexual reproduction carried out fully under water, including subaqueous pollination, locally clones can grow several football fields large (*36*). Previously, the accumulation of somatic genetic variation (as SNPs) has been observed in large mega-clones (*37*), but currently a dating of young, several year to decade old, clones is elusive, but highly useful to elucidate the demographic structure, longevity and ultimately vulnerability of seagrasses to a changing ocean environment.

To demonstrate that the epimutation-clock is a powerful tool for the reconstruction and dating of clonal phylogenies over short time scales, we took advantage of two *Z. marina* clones that had been cultured since 2004 for ecological experiments in Bodega Bay, California (**Materials and Methods, Supplementary text, Table S8-9**). These two clones (clone R and clone G) were initiated and propagated independently from each other in large tanks under ambient light, temperature and flow-through of ocean water (**Fig. 4A**). Sixteen ramets were sampled from each clone in 2021, corresponding to a clonal age of 17 years (**Materials and Methods**). WGBS was performed for 15 ramets of clone R, as one ramet did not belong to this clone (*38*). As for clone G, all 16 ramets were sequenced using WGBS, and technical replicates were generated by sequencing five aliquots of one sample independently (**Materials and Methods**). As extensive epigenomic information is lacking for the *de novo* identification of clock-like regions in *Z. marina*, we used gbM genes as a proxy of the epimutation-clock (**Materials and Methods**). For comparison, we also generated deep (100x) re-sequencing data for SNP identification in a subset of these ramets (**Materials and Methods**).

**Fig. 4.**
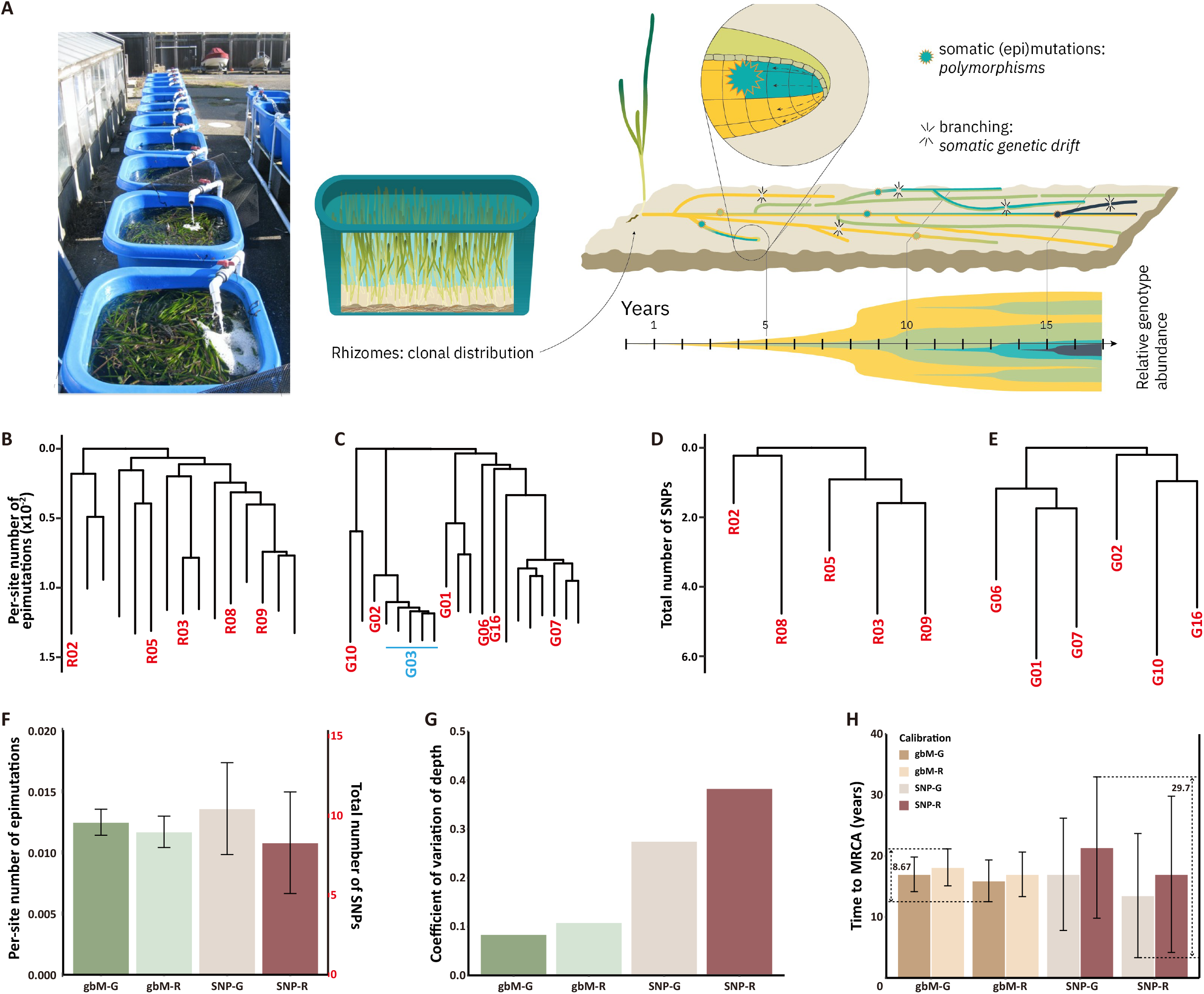
Evolutionary histories of *Z. marina* experimental clones. **(A)** Two independent *Z. marina* clonal lines were established from Bodega Harbor in large tanks under ambient light, temperature, and flow-through of ocean water. They were maintained via clonal reproduction for 17 years. **(B-C)** DNA methylomes were sequenced for 15 and 16 ramets of the R and G clones (five technique replicates were generated with the same ramet G03, blue). Using CG sites within gbM genes as a proxy of the epimutation clock, we inferred the phylogeny of the R and G clones separately. We re-sequenced genomes from five samples from the R clone and six samples from the G clone. Only 31 and 47 segregating SNPs were identified. We inferred the phylogeny using the segregating SNPs, and the trees of the R clone and the G clone are shown in **(D)** and **(E)**. **(F-G)**The average depth of two gbM-based trees is 1.246 x 10-2 and 1.168 x 10-2 with CVs less than 11%. However, the average depth in the SNP trees is much more variable, with CVs are over 27%. **(H)** The estimated times to MRCA were further calibrated with these depths and substitution rates.

Our analysis revealed that the clones have generated 20,713 (clone R) and 21,008 (clone G) fixed CG methylation changes in gbM genes over the course of 17 years, on average, compared with only 31 and 47 fixed SNP changes (**Materials and Methods**). Again, this constitutes a large excess of epimutations relative to SNPs per unit of time, even though the effective genome size of the former is orders of magnitude smaller. Although the true clonal phylogeny is unknown in this experiment, an epimutation-clock based analysis revealed high-confidence phylogenetic trees in both clones (**Fig. 4B-C, Supplementary text**). We were unable to recapitulate the phylogenetic trees using fixed SNPs (**Fig. 4D-E**). Instead, we observed that the bootstrap support values in the SNP-based trees were low (~0.6) (**Supplementary text**), which indicates that there is little phylogenetic information in the few SNPs captured among clonal ramets over these time scales.

To assess our ability to date the clones based on DNA methylation data, we calibrated our epimutation-clock in clone R, using its known age of exactly 17 years (**Method, Fig. 4 F-H**). Application of the calibrated clock to the phylogenetic tree of clone G estimates the MRCA at 18.14 ± 3.01 years ago (95% CI), which aligns well with the actual founding date of this clone. (**Fig. 4H, Table S9**). Applying the same approach using fixed SNPs led to variable estimates 21.39 ± 11.59 years ago (95% CI), an uncertainty of one decade over this short time-scales (**Fig. 4H**).

The existence of a fast-ticking evolutionary epigenetic clock in plants opens novel research avenues at the interface between evolutionary biology and ecology (*9*). Our proof-of-principle study focused on the application of this clock to the reconstruction and dating of shallow, intra-species phylogenies that arise within clonal or selfing species over short timescales. The use of epimutation-clocks may be extended to plant species with longer life-cycles as the necessary clock-calibration could rely on DNA methylation data from F2 intercrosses, rather than from mutation accumulation lines. A promising extension of our work is to combine the clock-like regions with modified coalescent methods to infer recent demographic changes (i.e. bottlenecks) and selection at the population level. The rapid rate at which epigenetic diversity arises within the clock-like regions will facilitate unprecedented temporal resolution. It can shed new light on microevolutionary questions that have been challenging to resolve, such as the timing of introduction of invasive species, the rate of poleward and upslope spread of species after the retreat of the Pleistocene glaciers, and the consequences of anthropogenic activities for population divergence. In an era of climate change, plant biodiversity is transforming at a fast pace. The ability to monitor rapid population dynamics at the molecular level will help us gauge the fate of this diversity going into the future.

## Acknowledgments

We would like to thank Susanne Landis (www.scienstration.com) for her help preparing the figures. TBHR acknowledges support from the HFSP project (PI TBHR; ADAPTASEX; RGP42/2020). RJS acknowledges support from the National Science Foundation (MCB-1856143) and the National Institutes of Health (R01GM134682). FJ acknowledges support from the Deutsche Forschungsgemeinschaft (DFG). Z.Z. and L.Y. hold fellowships from the China Scholarship Council (no. CSC202006380020) and (no. CSC201704910807), respectively.

